# *Aedes albopictus* is present in the lowlands of southern Zambia

**DOI:** 10.1101/2023.09.29.560125

**Authors:** Daniel R. Matute, Brandon S. Cooper

**Affiliations:** Biology Department, University of North Carolina, 250 Bell Tower Drive, Genome Sciences Building, Chapel Hill, NC 27510; Division of Biological Sciences, University of Montana, 32 Campus Dr., Missoula, MT 59812

## Abstract

Identifying the current geographic range of disease vectors is a critical first step towards determining effective mechanisms for controlling and potentially eradicating them. This is particularly true given that historical vector ranges may expand due to changing climates and human activity. The *Aedes* subgenus *Stegomyia* contains over 100 species, and among them, *Ae. aegypti* and *Ae. albopictus* mosquitoes represent the largest concern for public health, spreading dengue, chikungunya, and Zika viruses. While *Ae. aegypti* has been observed in the country of Zambia for decades, *Ae. albopictus* has not. In 2015 we sampled four urban and two rural areas in Zambia for *Aedes* species. Using DNA barcoding, we confirmed the presence of immature and adult *Ae. albopictus* at two rural sites: Siavonga and Livingstone. These genotypes seem most closely related to specimens previously collected in Mozambique based on CO1 sequence from mtDNA. We resampled Siavonga and Livingstone sites in 2019, again observing immature and adult *Ae. albopictus* at both sites. Relative *Ae. albopictus* frequencies were similar between sites, with the exception of immature life stages, which were higher in Siavonga than in Livingstone in 2019. While *Ae. albopictus* frequencies did not vary through time in Livingstone, both immature and adult frequencies increased through time in Siavonga. This report serves to document the presence of *Ae. albopictus* in Zambia, which will contribute to the process of determining the potential public health implications of this disease vector in Central Africa.

## INTRODUCTION

Diseases transmitted by Dipteran vectors represent one of the most important threats facing public health (Gubler 2009; Schmidt *et al*. 2013; Danasekaran *et al*. 2014; Torto and Tchouassi 2021). Approximately half of the world population may be at risk of contracting a vector-borne disease over their lifetime (Danasekaran *et al*. 2014; Torto and Tchouassi 2021), many of which are life-threatening and cause enormous cost to the patients and local public health systems they affect. The disease burden of Dengue, for example, seems to be largely unreported (Bhatt *et al*. 2013) and yet has risen dramatically over the last three decades (Bhatt *et al*. 2013; Du *et al*. 2021; Tian *et al*. 2022). Vectors are also important because they are effective conduits of zoonotic diseases that are a significant threat to human populations (Karesh *et al*. 2012; Schmidt *et al*. 2013; Gibb *et al*. 2020). The impact of vectors on public health is likely to increase as changing climates and anthropogenic activities influence species’ ranges and their contacts with human populations (Carlson *et al*. 2023; Longbottom *et al*. 2023).

Mosquitoes from the genus *Aedes* pose a significant threat to human health on multiple continents. *Aedes aegypti* is the primary vector of viral diseases such as dengue and yellow fever. Each year at least 50,000 patients die because of dengue (Wilder-Smith *et al*. 2010), and at least 30,000 because of yellow fever (Barnett 2007). These numbers are likely underestimated (Rigau-Pérez 2006; Barnett 2007; Allwinn *et al*. 2008; Adalja *et al*. 2012; Beaumier *et al*. 2014). *Aedes aegypti* is endemic to West Africa and has spread to more than 120 countries distributed across the globe (Rose *et al*. 2023). It has been hypothesized that *Ae. aegypti* range expansion has occurred for at least the last 500 years (Christophers 1960; Lounibos 2002)—this expansion may be as fast as 250 km per year in North America (Kraemer *et al*. 2019). A second species, *Ae. albopictus*, is endemic to Southeast Asia and has spread to all continents in the last five decades (Hawley 1988; BENEDICT *et al*. 2007; Paupy *et al*. 2009). In places like North America and Europe, the *Ae. albopictus* may be spreading over 100 km per year (Kraemer *et al*. 2019). Both species are expected to expand their geographic ranges as global temperatures increase (Fischer *et al*. 2014; Kamal *et al*. 2018; Kraemer *et al*. 2019; Liu *et al*. 2019; Laporta *et al*. 2023). This expansion is likely to impact human populations that have typically not been impacted by primarily tropical diseases.

In areas where vector-borne diseases are endemic, the invasion of *Ae. albopictus* is of particular importance for public health. Even though *Ae. albopictus* has historically been considered a secondary vector of dengue and yellow fever (reviewed in (Paupy *et al*. 2009)), recent work has demonstrated that *Ae. albopictus* is a competent vector of both diseases (Gloria-Soria *et al*. 2020). *Ae. albopictus* is also a vector of Zika and Chikungunya (Vega-Rúa *et al*. 2014; Garcia-Luna *et al*. 2018; McKenzie *et al*. 2019; Gloria-Soria *et al*. 2020), two emergent viral diseases with notable disease burden (Puntasecca *et al*. 2021). For example, in Cameroon and Gabón, the presence of *Ae. albopictus* was associated with epidemics of these two diseases in 2007 (Grard 2014). Both *Ae. aegypti* and *Ae. albopictus* are also potential vectors of other arboviruses that include Japanese encephalitis virus (Rosen *et al*. 1985), West Nile virus (Zhang *et al*. 2022), eastern equine encephalitis virus (Scott *et al*. 1990; Mitchell *et al*. 1992; Turell *et al*. 1994), and La Cross Virus (Gerhardt *et al*. 2001; Jackson *et al*. 2014; Westby *et al*. 2015). Thus, understanding the geographic distribution of these vectors remains critical.

*Aedes albopictus* is now prevalent across remote areas of Central Africa (Djiappi-Tchamen *et al*. 2021; Montgomery *et al*. 2022; Canelas *et al*. 2023; Obame-Nkoghe *et al*. 2023). The expansion, thus, has the potential to induce strain in health systems that are already overburdened. However, it is possible that *Ae. albopictus* might outcompete *Ae. aegypti*, through competitive exclusion ((O’meara *et al*. 1995; Britch *et al*. 2008; Juliano 2010) but see (Chan *et al*. 1971; Parker *et al*. 2019)), or satyrization (Zhou *et al*. 2022), which ultimately might reduce the population sizes of the more effective vector, *Ae. aegypti*. Understanding the precise geographic range of *Ae. albopictus* is an important public health issue that can answer the relative importance of the species as a disease vector and as a competitor in the vector species community, but one that can only be addressed with meticulous field collections.

Zambia is one of the few countries on the African continent where *Ae. Albopictus* has not been formally observed (Gubler 2003; Longbottom *et al*. 2023). Based on its wider distribution in Africa, we hypothesized that *Ae. albopictus* is likely present in Zambia. To test this, we sampled the *Aedes* of seven locations in Zambia in 2015, and two strategically chosen sites in 2019. We discovered both immature and adult *Ae. albopictus* individuals at two sites in southern Zambia and in both years, with *Ae. albopictus* frequency increasing through time in one. Based on this temporal and spatial presence of *Ae. albopictus*, we suggest that *Ae. albopictus* is a resident species in the lowlands of Zambia. We report that *Ae. albopictus* may be generally increasing in frequency, but this increase is unlikely to be rapid enough to displace *Ae. aegypti* in the near future. The isolates we observe are most closely related to isolates from Mozambique based on mtDNA barcoding. Our results will likely contribute to local control strategies, as the two species show differences in their preferred habitats.

## METHODS

### Locations

We sampled *Aedes* in seven different locations in 2015. This included sampling the boundaries of four major Zambian cities: Lusaka, Livingstone, Chipata, and Mazabuka. We also sampled three rural areas, including near Luangwa National Park, Siavonga (on the shores of Lake Kariba), and Shesheke. The 2019 sampling was strategically restricted to only two of these sites based on our 2015 observations: Livingstone and Siavonga. Table 1 lists the coordinates of the collection sites and the number of days that we sampled them.

**TABLE 1.** Coordinates and sampling outcomes of the 2015 and 2019 sampling sites. NA indicates that the site was not sampled in 2019. Imm: Immatures.

|  |  |  | 2015 |  |  | 2019 |  |  |
| --- | --- | --- | --- | --- | --- | --- | --- | --- |
|  | Altitude<br>(m) | Coordinates | Days | Imm. | Adults | Days | Imm. |  |
| Lusaka (C) | 1,274.9 | -15.504919,<br>28.265470 | 4 | 0 | 0 | NA | NA | NA |
| North<br>Luangwa<br>national<br>park (O) | 595.1 | -12.060373,<br>32.356497 | 5 | 0 | 0 | NA | NA | NA |
| Chipata<br>(C) | 1,184.7 | -13.621321,<br>32.652523 | 5 | 0 | 0 | NA | NA | NA |
| Mazabuka<br>(C) | 1,045.6 | -15.861312,<br>27.712943 | 3 | 0 | 0 | NA | NA | NA |
| Sesheke<br>(O) | 952.3 | -17.484532,<br>24.331769 | 5 | 0 | 0 | NA | NA | NA |
| Siavonga<br>(O) | 509.2 m | -16.542560,<br>28.689792 | 5 | 1 | 2 | 5 | 15 | 14 |
| Livingston<br>e (C) | 903.7 m | -17.871160,<br>25.857488 | 5 | 1 | 5 | 5 | 3 | 7 |

### Insect collections

Our goal was to determine the presence and frequency of immature and adult *Ae. albopictus* at the sites we sampled. To sample immatures, we collected larvae and pupae (collectively referred to as immature stages) by placing 32oz Ziploc containers (Reference number: 70937) with fish food and water, outside buildings (ovitraps). We then waited for an average of 3 to 5 days (Table 1), checking daily to make sure that animals did not disturb the traps. Immature stages were collected for ten days using a plastic pipette and placed in individual borosilicate vials. Each specimen was then inspected for taxonomically informative traits (See below).

To collect adults, we used CDC-type light traps (2836BQX, BioQuip; Rancho Domingo, CA) connected to a 6V as a power source. We used a mixture of yeast and water as a source of CO_2_. The trap was operated for three days at a time, but the collection sack was removed and the 6V battery was replaced every 48 hours. All the insects collected in the trap were removed from the traps using a mouth aspirator (1135A Aspirator–BioQuip; Rancho Domingo, CA), and anesthetized with FlyNap (Carolina Biological Supply Company, Burlington, NC). The totality of the collection was placed in a Petri dish and inspected under a Leica dissecting scope.

### Species identification

We differentiated between *Ae. albopictus* and *Ae. aegypti* larvae using a non-destructive procedure and two diagnostic traits. In *Ae. aegypti*, the comb scale on the terminal segment has a single row of pitchfork-shaped scales; larvae also have strong black hooks in the side of the thorax. In *Ae. albopictus*, the comb scale in the terminal segment is also in one row, but the scales are thorn-shaped and the hooks on the side of the thorax are small or absent (Lee 1998). Pupae were differentiated by the shape and length of the terminal setae in the paddle. While *Ae. aegypti* has short and simple terminal setae at the edge of the paddle, *Ae. albopictus* has long hairs (Penn 1949).

Larvae and pupae were kept in glass vials until they hatched and were inspected for adult traits that differentiate them. In *Ae. albopictus* the scutum has one silvery-white stripe down its middle, the thorax has patches of silver scales and a black clipeus. In *Ae. aegypti*, the scutum has a lyre-shaped pattern of white scales, a clipeus with white scales. Our field designations were then confirmed using genetic data.

### MtDNA typing

To confirm the accuracy of our field identifications using morphological traits, we used a mtDNA marker, cytochrome oxidase I (*COI*) to type species. Previous studies have shown that COI-based DNA barcoding is able to differentiate between species of *Aedes*, and other mosquito genera (Chan *et al*. 2014). We extracted DNA from single mosquitoes using the Qiagen DNeasy Blood and Tissue Kit (Qiagen Inc., Valencia, CA). We then amplified ∼25 ng of total DNA using PCR and conditions previously reported (Chan *et al*. 2014), using the two following primers: 5’-AAAAAGATGTATTTAAATTTCGGTCTG-3’ and 5’-TGTAATTGTTACTGCTCATGCTTTT-3’. Briefly, the amplification cycles were as follows: 94°C for 1 min followed by (34 cycles at 94°C for 30 seconds, annealing temperature: 52°C for 30 seconds, 72°C for 1 min 30 seconds), and a single final step at 72°C for 7 min. All amplifications were performed using a 2720 Thermal Cycler from Applied Biosystems (Foster City, CA).

PCR products were sent for sequencing to Eton Biosciences (NC) using the same PCR primers. Sequencing of PCR fragments was performed using the cyclic reaction termination method (Sanger sequencing) using the BigDye Terminator Cycle Sequencing Kits (Applied Biosystems, Foster City, CA, USA). To minimize false polymorphism, we sequenced both DNA strands, aligned them, and examined them with 4Peaks 1.7.1 (http://nucleobytes.com/index.php/4peaks). We extracted DNA from 31 adult individuals, and successfully PCR-amplified 24 samples (all samples collected in 2023) and obtained COI sequences from 18 amplicons. The resulting two haplotypes (See Results and Discussion) were deposited in Genbank (Accession numbers: TBD).

We obtained previously published sequences of *Ae. albopictus* to infer a crude genealogy of the species (Table S1), and to determine the most closely related geographic samples to our collections from Zambia. We used this COI alignment to generate a maximum-likelihood phylogenetic tree with IQTREE (Nguyen *et al*. 2015; Minh *et al*. 2020), using automatic inference of the model of molecular evolution (option -m), and a random seed (752482). We rooted the tree using *Ae. aegypti* (voucher ZOOMENTAa8, genBank accession number: MK265729). We used ultrafast bootstrap to calculate the branch support of each node (Minh *et al*. 2013; Hoang *et al*. 2018)

## RESULTS AND DISCUSSION

Our sampling revealed the presence of *Ae. albopictus* in two out of the seven locations that we sampled in Zambia: Siavonga (509m) and Livingstone (903m). Environmental niche models have previously predicted Siavonga as a favorable site for *Aedes* occurrence (Velu *et al*. 2021). *Aedes albopictus* was absent from collections we made at the three urban locations with higher altitude (Lusaka: 1,275; Mazabuka: 1,046m; and Chipata: 1,185m) and from two of the other rural locations (North Luangwa National Park: 595m, and Shesheke: 952m).

Next, we studied the frequency of *Ae. albopictus* relative to *Ae. aegypti* in Siavonga and Livingstone. In 2015, the relative proportion of immature *Ae. albopictus* individuals was similar in Siavonga (1.5%) and Livingstone (2%; χ ^2^ < 0.01, df = 1, *P* = 1). The relative proportion of *Ae. albopictus* adults also did not differ significantly between sites (Siavonga = 3.1%, Livingstone = 8.3%, χ ^2^ = 0.788, df = 1, *P* = 0.375). In 2019, immature stages of *Ae. albopictus* were proportionally more abundant in Siavonga (14%) than in Livingstone (3%, χ ^2^ = 8.03, df = 1, *P* = 4.6 × 10^−3^), but again there was no significant difference in the relative frequency of *Ae. albopictus* adults between the two locations (Siavonga: 18.4%, Livingstone: 8%, χ ^2^ = 3.119, df = 1, *P* = 0.077). These results motivate a large-scale and longitudinal study of *Ae. albopictus* abundance in the lowlands of Zambia.

We also studied whether the relative proportion of *Ae. albopictus* varied between years in Siavonga and Livingstone. Between 2015 and 2019, the proportion of *Ae. albopictus* immature individuals increased from 1.5% to 14% in Siavonga (χ ^2^ = 6.713, df = 1, *P* = 9.573 × 10^−3^) but the relative proportion of *Ae. albopictus* immature individuals did not not change between years in Livingstone (2-3%;^χ 2^ < 0.01, df = 1, *P* = 0.95). We observed a similar result for adults, where adult *Ae. albopictus* became more common in Siavonga, increasing in frequency from 3.1% to 18.4% between 2015 and 2019 (χ ^2^ = 6.745, df = 1, *P* = 9.40 × 10^−3^); but the relative proportion of *Ae. albopictus* adults remained similar between years in Livingstone (8.3%-8.0, χ ^2^ < 0.01, df = 1, *P*= 1).

The absence of *Ae. albopictus* from locations with an elevation over 1,000 meters is also noteworthy. While several environmental niche modeling efforts have suggested that *Ae. albopictus* may have the capacity to expand its range into higher altitudes (Fischer *et al*. 2014; Echeverry-Cárdenas *et al*. 2021; Santos *et al*. 2022), longitudinal sampling of mountainous regions in Zambia and other areas of Africa is needed to test this hypothesis. Sampling of other continents supports *Ae. albopictus* expansion into higher altitudes. For example, *Ae. albopictus* occurs at significantly higher altitudes in South America than the median altitude in Zambia (∼1,200m). In Colombia, *Ae. albopictus* is routinely sampled at altitudes similar to Lusaka in Zambia (up to 1,300m, (Echeverry-Cárdenas *et al*. 2021)). In a more extreme case, *Ae. albopictus* was recently sampled in Bolivia at an elevation of over 2,300m (Ríos *et al*. 2023). Together, these observations suggest that while altitude may influence rates of *Ae. albopictus*’ dispersal, it is not an insurmountable barrier for its establishment.

We next used COI sequence from mtDNA to infer the general genetic relationships among the specimens of *Ae. albopictus* we sampled from Zambia and isolates collected in other countries. Zambia is a landlocked country, but it shares borders with countries in the Indian Ocean. This proximity is relevant because the first observations of *Ae. albopictus* in Africa were from samples collected from Indian Ocean islands (Madagascar: 1904, Mauritius: 1922, Seychelles: 1977, reviewed in (Longbottom *et al*. 2023)). We find that the isolates of *Ae. albopictus* we sampled from Siavonga, Zambia are closely related to other mtDNA haplotypes present in Mozambique, suggesting that the invasion in Zambia likely occurred via migration from adjacent countries. Notably, we observe two different *COI Ae. albopictus* haplotypes, both of which are present in Mozambique. Of the 18 individuals that we typed using COI sequence, 12 have a *COI* haplotype identical to a sample collected in Mozambique (LC726293.1), and six have a haplotype identical to a second Mozambique specimen (LC726392.1). While DNA barcoding is powerful for species identification, the typing here is inherently limited for the identification of demographic events. The inference of the events and population of origin that lead to the presence of *Ae. albopictus* in Zambia will require more systematic insect sampling and sequencing of whole genomes.

While we do not currently know the full distribution of *Ae. albopictus* in Zambia, or across Africa more broadly, our observation of *Ae. albopictus* at two sites in two years significantly contribute to current knowledge by establishing that by 2015 *Ae. albopictus* colonized the southern lowlands of Zambia (under 1,000 m above the sea level); and in the case of Siavonga, our results indicate that the relative frequency of *Ae. albopictus* is increasing through time. Notably, surrounding counties such as Mozambique (Kampango and Abílio 2016; Abílio *et al*. 2018), Tanzania (Patrick *et al*. 2017), and the Democratic Republic of the Congo (Bobanga *et al*. 2018; Vulu *et al*. 2021) had reported the presence of *Ae. albopictus* at a similar time (2015-2016). This may indicate recent expansion of *Ae. albopictus* into these countries and Zambia. In contrast, it is plausible that *Ae. albopictus* has gone undetected, and successful independent sampling of each country randomly occurred at a similar time.

The finding of *Ae. albopictus* in Zambia is of importance for public health because several diseases transmitted by *Aedes* species—including Dengue, yellow fever, and Chikunguya—are present in Zambia. For example, a serological survey revealed that 4.1% of participants from Western and North-Western Zambia were seropositive for Dengue (*N* = 3,624, (Mazaba-Liwewe *et al*. 2014)). In central Zambia, the proportion of Dengue seropositive patients seems to be higher (16.8%, *N* = 214; (Chisenga *et al*. 2020)). The same survey found that 36.9% of patients from Central Zambia were seropositive for Chikungunya, 10.8% for Zika, and 19.6% for Mayaro (Chisenga *et al*. 2020). To our knowledge, no serological prevalence surveys for any of these diseases exist for areas where we have sampled *Ae. albopictus*. However, there is evidence that Zika exists in this region, with twenty samples out of 50 from non-human primate samples (*Chlorocebus cynosuros* and *Papio ursinus*, (Wastika *et al*. 2019)) having anti-Zika antibodies. This suggests that non-human primates might serve as Zika virus reservoirs in areas where we sampled *Ae. albopictus*. Other diseases are also worth considering. For example, West Nile virus has also been isolated from other mosquitoes (Orba *et al*. 2018), humans (Mweene-Ndumba *et al*. 2015) and potential reservoir species (Simulundu *et al*. 2020); and *Ae. albopictus* is an effective vector of West Nile virus (Zhang *et al*. 2022). The confluence of these vectors and pathogens they may plausibly transmit motivates future analyses focused on better understanding disease prevalence across Zambia and other regions of Africa.

## ACKNOWLEDGEMENTS

We would like to thank the Matute lab for helpful comments. DRM was supported by the National Institute of General Medical Sciences under Award R35GM148244. BSC was supported by the National Institute of General Medical Sciences under Award R35GM124701.

## FIGURES

**FIGURE 1.**
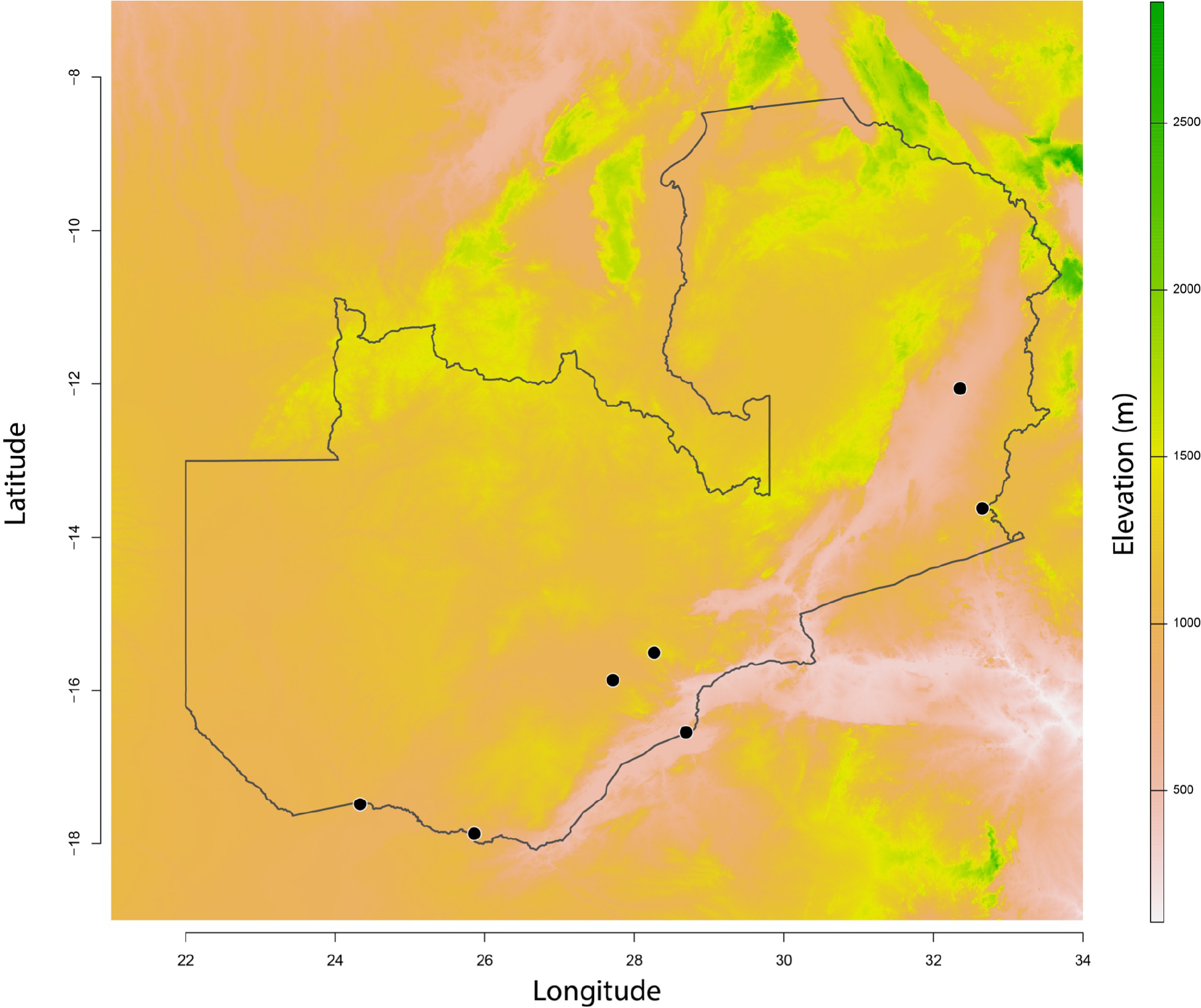
Sites included in this study. We collected seven different sites, four of which were at the boundaries of urban centers (C) and three of which were in rural areas. Two sites, Siavonga and Livingstone (marked with a star) were sampled in 2015 and 2019.

**FIGURE 2.**
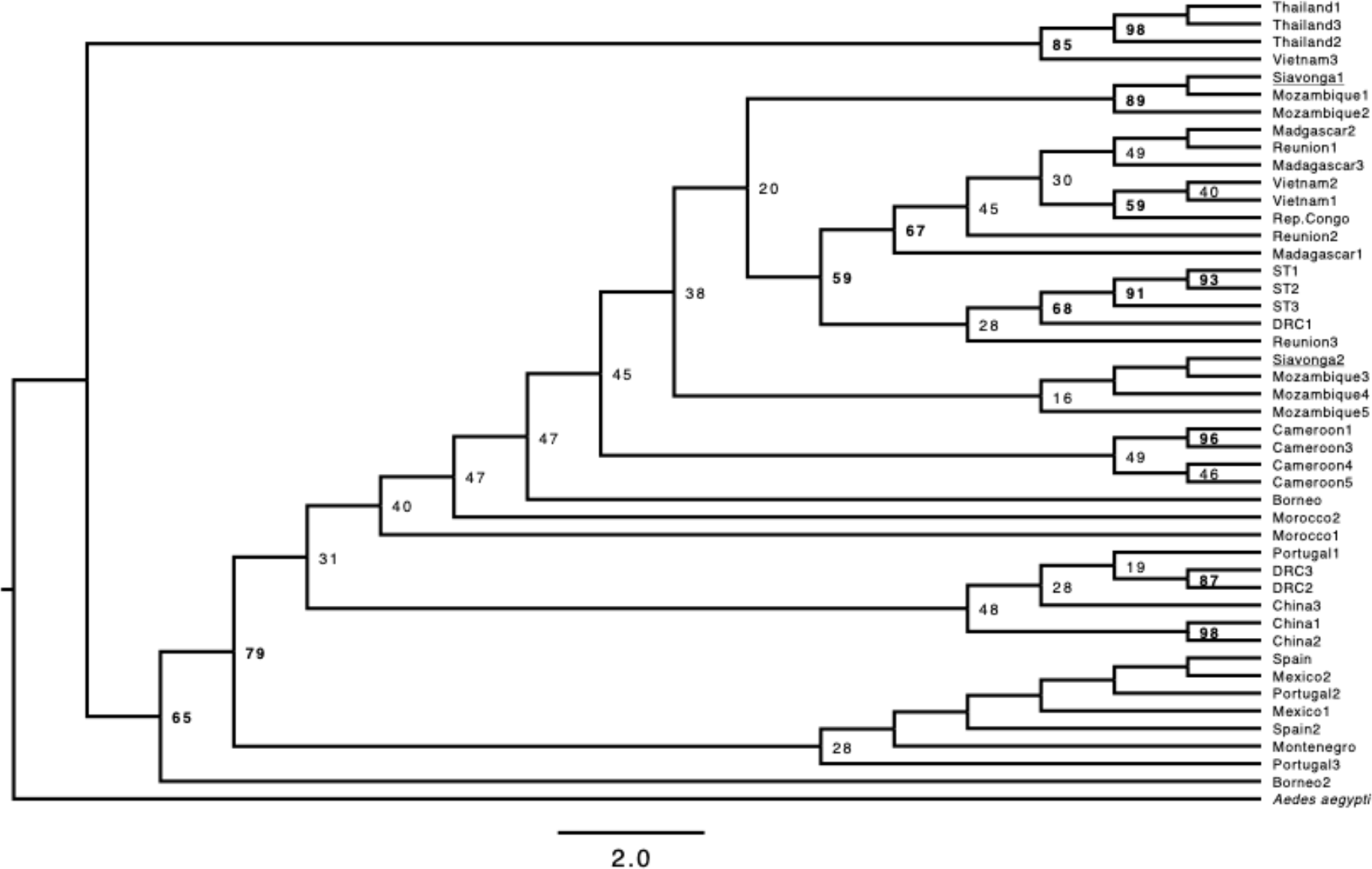
Maximum likelihood tree derived from the COI mtDNA barcode. We retained all the branches in spite of some low-support branches. Values above each node correspond to the bootstrap support. Depicted branch lengths are not proportional to facilitate the interpretation of the genealogical relationships. Figure S1 shows a scaled tree.

