## Supplementary material for "*Aedes albopictus* is present in the lowlands of southern Zambia": Table S1, Figure S1

**TABLE S1.** Genbank accession numbers for the COI barcodes used to generate an *Ae. albopictus* gene genealogy.

| **Code in Figures 2 and S1** | **Gene Bank accession number** |
|---|---|
| *Aedes aegypti* | MK265729 |
| Thailand1 | KM613129.1 |
| Thailand3 | KM613121.1 |
| Thailand2 | KM613128.1 |
| Vietnam3 | HQ398902.1 |
| Siavonga1 | TBD |
| Mozambique1 | LC726387.1 |
| Mozambique2 | LC726393.1 |
| Siavonga2 | TBD |
| Mozambique3 | LC726394.1 |
| Mozambique4 | LC726392.1 |
| Mozambique5 | LC726380.1 |
| Borneo | MN540323.1 |
| Borneo2 | MN540322.1 |
| Spain | KU319448.1 |
| Portugal3 | MK995330.1 |
| Portugal2 | MK995331.1 |
| Mexico1 | MT999274.1 |

| | |
|---|---|
| Mexico2 | MT552470.1 |
| Montenegro | MK505589.1 |
| Spain2 | KU319443.1 |
| Portugal1 | MK995332 |
| DRC3 | MT345383 |
| Morocco2 | KU522419 |
| Morocco1 | KU522421 |
| DRC2 | MT345388 |
| China3 | KX886337 |
| China1 | KX981869 |
| China2 | KX981868 |
| Cameroon1 | MH921572 |
| Cameroon3 | MH921568 |
| Cameroon4 | MH921571 |
| Cameroon2 | MH921570 |
| Madagascar2 | JN406732 |
| Madagascar3 | JN406725 |
| Reunion1 | JN406663 |
| Vietnam2 | KX573911 |
| Vietnam1 | KX495925 |
| ST1 | JF309319 |
| ST2 | JF309319 |
| ST3 | JF309318 |
| DRC1 | MT345390 |
| Reunion2 | AJ971012 |
| Rep. Congo | MH025948 |

| | |
|---|---|
| Madagascar1 | AJ971007 |
| Reunion2 | AJ971013 |

**FIGURE S1. Maximum likelihood *Ae. albopictus* tree derived from the *COI* mtDNA barcode.** We retained all the branches in spite of some low-support branches. Values above each node correspond to the bootstrap support. Branch lengths are proportional but the topology is identical to the one shown in Figure 2.

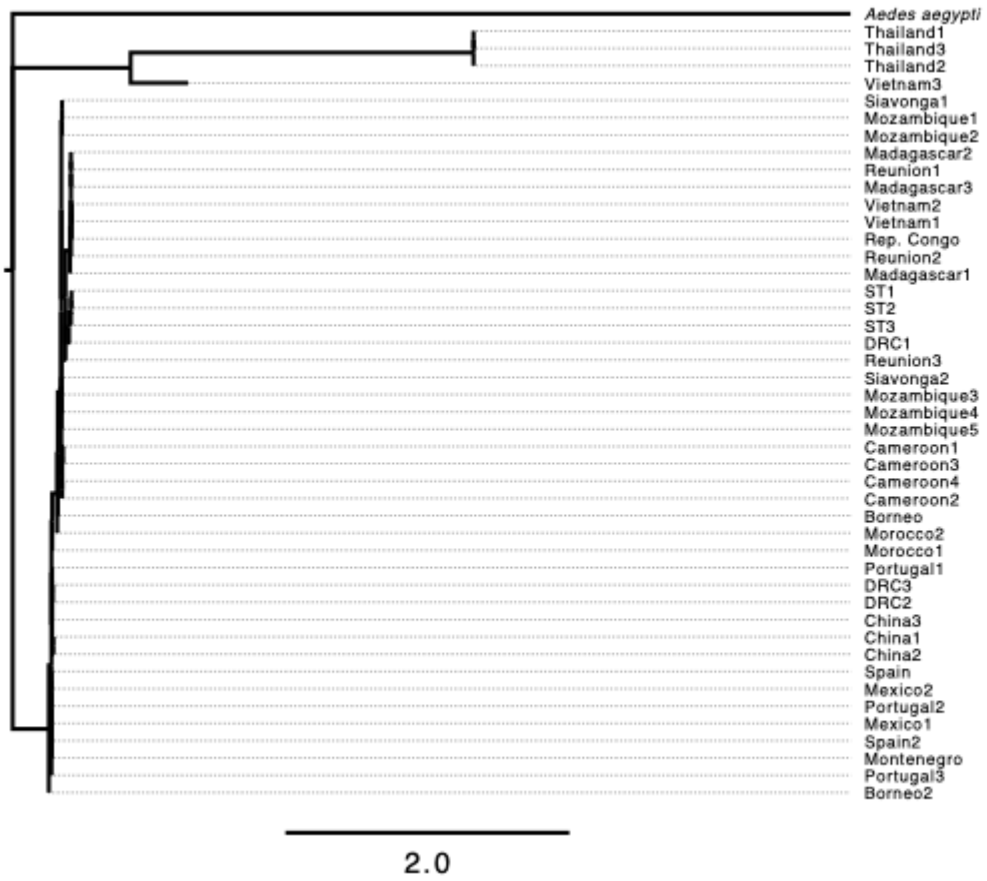
